# Beyond fish eDNA metabarcoding: Field replicates disproportionately improve the detection of stream associated vertebrate species

**DOI:** 10.1101/2021.03.26.437227

**Authors:** Till-Hendrik Macher, Robin Schütz, Jens Arle, Arne J. Beermann, Jan Koschorreck, Florian Leese

## Abstract

Fast, reliable, and comprehensive biodiversity monitoring data are needed for environmental decision making and management. Recent work on fish environmental DNA (eDNA) metabarcoding shows that aquatic diversity can be captured fast, reliably, and non-invasively at moderate costs. Because water in a catchment flows to the lowest point in the landscape, often a stream, it can often collect traces of terrestrial species via surface or subsurface runoff along its way or when specimens come into direct contact with water (e.g., for drinking purposes). Thus, fish eDNA metabarcoding data can provide information on fish but also on other vertebrate species that live in riparian habitats. This additional data may offer a much more comprehensive approach for assessing vertebrate diversity at no additional costs. Studies on how the sampling strategy affects species detection especially of stream-associated communities, however, are scarce. We therefore performed an analysis on the effects of biological replication on both fish as well as (semi-)terrestrial species detection. Along a 2 km stretch of the river Mulde (Germany), we collected 18 1-L water samples and analyzed the relation of detected species richness and quantity of biological replicates taken. We detected 58 vertebrate species, of which 25 were fish and lamprey, 18 mammals, and 15 birds, which account for 50%, 24%, and 7% of all native species to the German federal state of Saxony-Anhalt. However, while increasing the number of biological replicates resulted in only 25% more detected fish and lamprey species, mammal, and bird species richness increased disproportionately by 69% and 84%, respectively. Contrary, PCR replicates showed little stochasticity. We thus emphasize to increase the number of biological replicates when the aim is to improve general species detections. This holds especially true, when the focus is on rare aquatic taxa or on (semi-)terrestrial species, the so-called ‘bycatch’. As a clear advantage, this information can be obtained without any additional sampling or laboratory effort when the sampling strategy is chosen carefully. With the increased use of eDNA metabarcoding as part of national fish bioassessment and monitoring programs, the complimentary information provided on bycatch can be used for biodiversity monitoring and conservation on a much broader scale.

## Introduction

Environmental DNA (eDNA) metabarcoding is a powerful and nowadays frequently applied method to assess and monitor fish biodiversity in streams (Cantera et al. 2019), lakes (Muri et al. 2020) and the sea (Andruszkiewicz et al. 2017). Contrary to conventional methods, such as net trapping or electrofishing, eDNA metabarcoding from water samples is non-invasive, safe and simple, and taxonomic richness estimates are generally more complete than classical assessments (Bernd Hänfling et al. 2016; Pont et al. 2018; Boivin‐Delisle et al. 2021). In view of the maturity of the method, the uptake of fish eDNA metabarcoding into regulatory monitoring programs, such as the European Water Framework Directive (2000/60/EC, WFD), is discussed (Hering et al. 2018; Pont et al. 2021).

In view of global biodiversity loss and the demand for spatio-temporally highly resolved data, eDNA metabarcoding has an additional, so far less explored potential: While fish species are primary targets, eDNA monitoring data can also provide reliable information on many other taxa either living in or in the vicinity of water bodies such as mammals (Andruszkiewicz et al. 2017; Closek et al. 2019), amphibians (Bálint et al. 2018; Lacoursière-Roussel et al. 2016; Harper et al. 2018), and birds (Ushio, Murata, et al. 2018; Day et al. 2019; Schütz, Tollrian, and Schweinsberg 2020). While traditional monitoring of birds is usually conducted by many hobby and professional ornithologists, the monitoring of mammals relies on far more advanced, non-invasive, observational methods such as camera traps or identification of field traces (e.g., hair or feces). Nevertheless, semi-aquatic, terrestrial, and aerial species emit genetic material to their environment, which allows their identification by eDNA-based approaches. These bycatches from one monitoring approach, as e.g., fish eDNA metabarcoding from water samples, can become important sources for other regulatory frameworks: While birds and mammals are not considered in the WFD, they are subject to the EU birds directive (Directive 2009/147/EC, 2009), the “EU Regulation 1143/2014 on Invasive Alien Species”, and the EU habitats directive (Council Directive 92/43/EEC, 1992). Monitoring data on birds and mammals are furthermore of major interest under the convention on biological diversity (see https://www.cbd.int/) and may become increasingly the basis of inventory estimates for regional, national, and international red lists (e.g., IUCN). The definition of bycatch and target, respectively, is artificially defined by the respective national or international regulations and directives. This differentiation of bycatch and target is irrelevant on the molecular level of eDNA, since eDNA from all different groups can be found in a single water sample. Thus, eDNA metabarcoding allows insights into the whole stream associated vertebrate community (Deiner et al. 2017; Ushio et al. 2017; Mariani et al. 2021), detecting not only aquatic but also semi-aquatic and terrestrial mammals and birds (figure 1). The collection of eDNA samples during monitoring studies thus can provide highly valuable information of a much broader scale without any (if the same metabarcoding primers are used) additional costs or sampling effort. Often universal, i.e., degenerate primers (Riaz et al. 2011; Miya et al. 2015; Taberlet et al. 2018), have the potential to efficiently target fish and lamprey, and moreover also to amplify DNA of species of birds and mammals as a bycatch, without reducing the fish detection rate.

**Figure 1:**
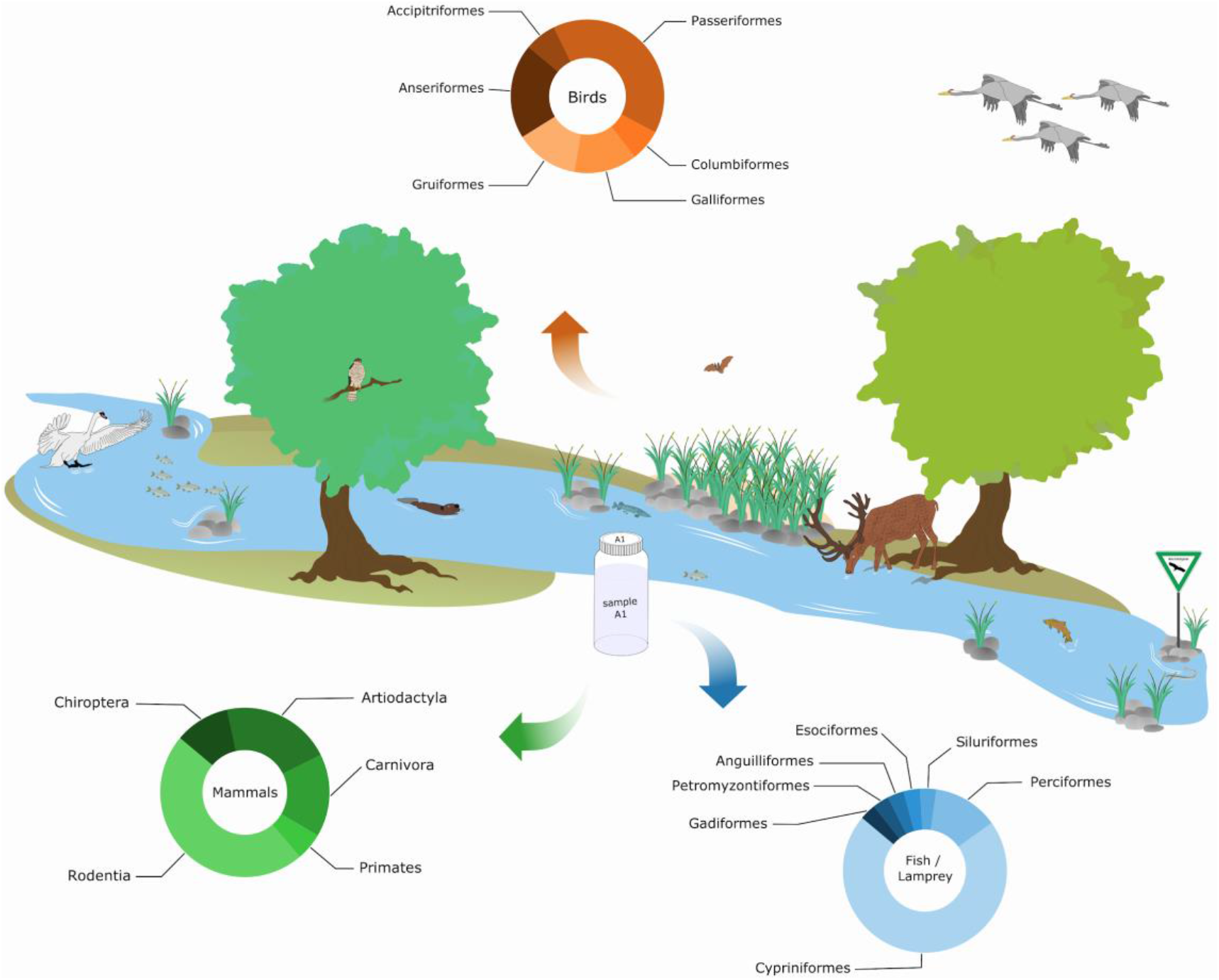
Scheme of a freshwater associated vertebrate community including some of the detected species. The OTU distribution among the classes of birds, mammals and fish/lamprey found in this study are illustrated in pie charts.

While the view of water bodies as ‘sinks’ or ‘conveyor belts’ (sensu Deiner et al. 2016) is appealing in view of holistic biodiversity monitoring, several issues are obvious. Typically, the non-target (semi-) terrestrial bycatch is difficult to detect. Especially when eDNA is not homogeneously distributed (Furlan et al. 2016; Cantera et al. 2019; Jeunen et al. 2019). Previous studies reported that the number of sampling sites and biological replicates can strongly influence the detected species richness (Civade et al. 2016; Bernd Hänfling et al. 2016; Valentini et al. 2016; Evans et al. 2017; Bálint et al. 2018; Doble et al. 2020). This holds true in particular for standing water bodies with strong stratification (Jeunen et al. 2019). For stream ecosystems, however, eDNA distribution can be assumed to be more homogeneous given turbulent flow. However, only a few studies tested this. For example, in a study by Cantera et al. (2019) tropical fish richness estimates showed that the filtration of 34 to 68 liters was sufficient to inventory the local fish fauna, while the filtration of larger volumes only slightly increased the detected species richness. However, this study focused on total fish diversity and did not consider other taxa. In addition, an important consideration from the practical standpoint of routine biomonitoring are trade-offs between sample number or water volume filtered and the actual increase in species detection with more samples or higher volumes. Given limited resources and time, the best compromise between sample number and detection probability is needed.

Therefore, we performed an eDNA metabarcoding survey using universal fish primers on water samples collected from the German river Mulde to assess the fish and stream associated vertebrate (bycatch) community. Our aims were i) to test the effect of biological sample replication on the detected fish species richness, and ii) to investigate the detection rate of usually discarded bycatch vertebrate species.

## Methods

### Sampling site

The sampling site was located at the Mulde weir in Dessau (Germany, 51°49’56.2“N 12°15’05.1“E). The river Mulde is a tributary of the Elbe system with an average effluent at the sampling site of 62.7 m^3^/s in April (2012-2018; (FIS FGG Elbe). From the complete stream system up to 34 fish species are reported (Geisler 2001, MULE fish report 2014), which is close to the total number of 50 fish species reported for the German federal state of Saxony-Anhalt (Kammerad et al. 2020). Amongst these are endangered and strictly protected fish as well as diadromous and invasive species. In accordance with the EU Water Framework Directive, a fish ladder was built in 2017 to surpass the 2.4 m weir and to allow for unimpeded migration of organisms, in particular fish.

### eDNA sampling

We collected 18 water samples in April 2019 over a stretch of 2 km: 4 samples each were collected 1 km upstream of the weir (location S1), directly upstream (S2) and directly downstream of the fish ladder (S3), and 1 km downstream of the weir (S4). Additionally, two samples were taken directly in the fish ladder itself (L1). For each sample, 1-L of water was collected in a sterile plastic bottle. To prevent cross-contamination, sterile laboratory gloves were changed between samples. All water samples were filtered on site to avoid contaminations and ease the transportation. Open MCE (mixed cellulose ester membrane) filters with a 0.45 µm pore size (diameter 47 mm, Nalgene) were used for the filtration. The filters were handled with sterile forceps and gloves were changed between each sample. An electric vacuum pump, a funnel filter, and a filter flask were installed for filtering the water. As field blanks, a total of two blank filters were placed on the filter flask and exposed to air for 20 seconds. The filters were transferred to 1.5 mL Eppendorf tubes filled with 96% ethanol, kept at 4°C overnight and then stored at −20°C until DNA extraction.

### DNA extraction

All laboratory steps were conducted under sterile conditions in a dedicated sterile laboratory (UV lights, sterile benches, overalls, gloves, and face masks). The filters were dried separately in sterile petri dishes and covered with aluminum foil overnight. Afterwards the filters were torn into pieces using sterile forceps and transferred into new 1.5 mL Eppendorf tubes. Subsequently, filters were eluted in 600 μL TNES-Buffer and 10 μL Proteinase K and incubated at 55°C and 1000 rpm for three hours. DNA was extracted from the filters following an adapted salt precipitation protocol (Weiss and Leese 2016) eluted in 50 μL PCR H_2_O and stored overnight at 4°C. Next, 0.5 μL RNase A (10mg/mL) was added to each sample and incubated for 30 minutes at 37°C on an Eppendorf ThermoMixer C (Eppendorf AG, Hamburg, Germany). Subsequently, samples were purified using the Qiagen MinElute DNeasy Blood & Tissue Kit (Hilden, Germany), following the manufacturer's protocol. Samples were eluted in 30 μL PCR-grade H_2_O.

### DNA amplification and sequencing

A two-step PCR approach was applied for amplifying the extracted DNA. In the first PCR, the vertebrate teleo2 primers (Taberlet et al. 2018) were used, that are optimized for European freshwater fish targeting a 129-209 bp long 12S gene fragment. In total, 100 first step PCR reactions were conducted, including 5 replicates per sample as well as 8 negative PCR controls and 2 field blanks. The PCR reaction volume was 50 μL consisting of 21 μL H_2_O, 25 μL Multiplex Mastermix (Qiagen Multiplex PCR Plus Kit, Qiagen, Hilden, Germany), 1 μL teleo02 forward primer and 1 μL teleo02 reverse primer and 2 μL of DNA template. The first PCR step was carried out at 95°C for 5 minutes followed by 35 cycles with 94°C for 30 seconds, 52°C for 90 seconds and 72°C for 90 seconds. The final elongation was carried out at 68°C for 10 minutes. After the first-step PCR, all five replicates of each sample were pooled together. For the second-step PCR, a universal tagging primer set was used (Buchner et al. in prep). A total of 52 second-step PCR reactions were conducted using two PCR replicates per sample, four first-step negative controls, four second-step negative controls and two field blanks. The PCR mix per sample contained 19 μL of H_2_O, 25 μL of Multiplex Mix, 2 μL combined primer (10 μM) and 4 μL first-step product. PCR conditions were 95°C for 5 minutes followed by 10 cycles at 94°C for 30 seconds, 62°C for 90 seconds and 72°C for 90 seconds. The final elongation was carried out at 68°C for 10 minutes. Following the second-step PCR, the PCR products were visualized on a 1% agarose gel to evaluate the amplification success. The samples were subsequently normalized to 25 ng per sample, using a SequalPrep Normalization Plate (Applied Biosystems, Foster City, CA, USA) following the manufacturer’s protocol. Subsequently, the normalized samples were pooled into one library. After library-pooling, the samples were concentrated using a NucleoSpin Gel and PCR Clean-up kit (Machery Nagel, Düren, Germany) following the manufacturer’s protocol. The final elution volume of the library was 22 μL. The samples were then analyzed using a Fragment Analyzer (High Sensitivity NGS Fragment Analysis Kit; Advanced Analytical, Ankeny, USA) to check for potential primer dimers and co-amplification and quantify the DNA concentration of the library. Primer dimers were removed by extracting PCR products using two lanes (10 μL each) of an E-Gel Power Snap Electrophoresis Device (ThermoFisher Scientific, Germany). This resulting library was sequenced on a MiSeq v2 250 bp PE Illumina at CeGaT (Tübingen, Germany).

### Bioinformatics

Raw reads for both libraries were received as demultiplexed fastq files. The quality of the raw reads was checked using FastQC (Andrews 2010). Paired-end reads were merged using VSEARCH version 2.11.1 (Rognes et al. 2016), allowing for 25% differences between merged pairs and a minimum overlap of 5 bp. Afterwards, primers were trimmed using cutadapt version 2.8 (Martin 2011). Reads were then filtered by length (119-219 bp threshold for teleo2 target fragment) and by maximum expected error (threshold below maxee = 1), using VSEARCH. The filtered reads were dereplicated and singletons and chimeras were removed with VSEARCH. All reads were then pooled using a custom python script and again dereplicated. Operational taxonomic units (OTUs) were obtained with a 97% similarity clustering and the seeding sequences were extracted as representative OTU sequences. The OTUs were remapped (usearch_global function, 97% similarity) to the individual sample files to create the read table. The read table was filtered by column (read abundance threshold: >0.01% of reads to keep the OTU) and then by row (OTU must be present in at least one of the samples). OTUs were blasted (web blast, blastn suite, nt database, blastn) against the Nation Center for Biotechnology Information (NCBI) database. The results were downloaded in xml format and processed using a custom python script (https://github.com/TillMacher/xml2_to_TTT). Here, the taxon ID and blast similarity was fetched from the xml file (suppl. table 1 sheet “Raw hits”) and the according taxonomy was downloaded from the NCBI server (suppl. table 1 “Taxonomy added”). The blast results were subsequently filtered in three steps. First, only the hit with the highest similarity was kept and duplicate hits were removed. When two or more different taxon names were found, all of them were kept. Subsequently, the hit table was filtered according to the thresholds described in JAMP (https://github.com/VascoElbrecht/JAMP), with a <97 % cutoff threshold to cut species level, <95 % for genus, <90 % for family, <85 % for order and below 85 % for class level (suppl. table 1 “JAMP filtering”). Subsequently, all remaining hits of one OTU were trimmed to their first shared taxonomic rank. Remaining duplicates (i.e., hits of one OTU that share the same taxonomy after the filtering) were dereplicated. Thus, each OTU was assigned to one taxonomic hit in the final taxonomy table (suppl. table 1 “JAMP hit”). Finally, OTUs were matched with the read table and OTUs that were not taxonomically assigned during the blast were discarded.

Both, the taxonomy and read table file were converted to the TaXon table format (suppl. table 2) for downstream analyses in TaxonTableTools v1.2.4 (Macher, Beermann, and Leese 2020). The separately sequenced PCR and field replicates were analyzed using the replicates analysis tools (correlation analyses and shared OTUs) and were subsequently merged. Furthermore, negative controls were excluded from the downstream dataset and only hits of the phylum Chordata were kept for the downstream analyses (suppl. table 3). PCR replicates were tested for correlations of number of reads and OTUs, based on Spearman's rank correlation coefficient. The read proportions, number of OTUs and number of unique species for each class were calculated. A Jaccard distance-based non-metric multidimensional scaling analysis (NMDS) was conducted to test if site effects between the five sampling locations were present, and all samples can be treated as individual field replicates (NMDS settings: 8000 iterations and 400 different initializations). Three rarefaction analyses were performed to calculate the effect of field replicates on the number of obtained species. Random sub-samples were drawn from all 18 field replicates and the number of observed fish/lamprey, bird and mammal species count were counted separately. Each draw was repeated 1000 times to account for stochastic effects. An occupancy plot was calculated to investigate the relative appearance of each species across all replicates. The plot was subsequently adjusted in Inkscape to add an order-specific color code.

## Results

We obtained 9,906,197 raw reads with 1,193,233 reads assigned to negative controls. After final quality filtering 7,520,725 reads remained (1,646 reads in negative controls), which were clustered into 474 OTUs. The sum of the reads in negative controls after clustering and remapping was 1,376. After the 0.01% threshold filtering of the read table, 153 OTUs remained of which we could assign 147 taxonomically. In five cases where the marker resolution was too low to distinguish between species, taxonomic annotation was manually edited to retain both species names. Therefore, we counted those cases as one entry in the species list since at least one is present (i.e., *Pipistrellus pipistrellus* / *P*. *pygmaeus*, *Blicca bjoerkna* / *Vimba vimba*, *Carassius auratus* / *C. carassius, Leuciscus aspius* / *Alburnus alburnus*). In an additional case OTU 17 was automatically reduced to genus level due to two 100% similarity reference sequences representing two different species, the European eel (*Anguilla anguilla*), and the American eel (*Anguilla rostrata*). Since the European eel is the only representative of its genus in Europe, we assigned the OTU manually to *Anguilla anguilla*. Furthermore, we assigned OTU 10 to the mallard (*Anas platyrhynchos*), after manually investigating the taxonomic assignment results. Due to various reference sequences of mallard breeds and one common shelduck breed (*Tadorna tadorna*), the automatic assignment was unable to find a consensus and thus reduced the taxonomic resolution to Anatidae level.

Three OTUs were assigned to Proteobacteria, Verrucomicrobia, and Bacteroidetes. These were removed for downstream analyses. The majority of reads in negative controls (1371) were found in one field negative control and were mostly assigned to *Sus scrofa*. Thus, the *Sus scrofa* OTU was excluded from the dataset. After merging replicates (OTUs that were not present in both replicates were discarded) and removal of negative controls, 137 vertebrate OTUs remained, 64 of which could be assigned to species level (suppl. table 3). Reads were mainly assigned to fish (Actinopterygii, 92% of all reads), while Hyperoartia (only recent representatives are lampreys) accounted for 0.1% of the reads. Mammals were represented by 6% of all reads and birds (Aves) by 2% (Figure 2B). Overall, 74 OTUs were assigned to fish, including 24 different species, while one OTU on species level was assigned to Hyperoartia. Furthermore, 17 OTUs were assigned to 15 bird species, and 44 OTUs to 18 different mammal species (figure 2B). The 25 fish and lamprey species (in the following summarized as fish/lamprey if not stated otherwise) belonged to the orders of Cypriniformes, Perciformes, Siluriformes, Esociformes, Anguilliformes, Petromyzontiformes, and Gadiformes (table 1). They account for 25 of 50 reported fish species from the German federal state Saxony-Anhalt (red list of Saxony-Anhalt, LAU 01/20). The overall 18 mammal species belonged to the orders of Rodentia, Primates, Carnivora, Artiodactyla, and Chiroptera (table 2), and the 15 bird species to Accipitriformes, Anseriformes, Gruiformes, Galliformes, Columbiformes, and Passeriformes (table 3). They account for 18 of 81 mammal species (22.2%) and 15 of 202 breeding bird species (7.4%) that are native to Saxony-Anhalt. In terms of read abundance the common dace (*Leuciscus leuciscus*) was the most abundant chordate species with 58% of all reads. Three further fish species showed read proportions of more than 2%, i.e., *Gymnocephalus cernua* (4%), *Abramis brama* (4%), and *Rutilus rutilus* (3%). The only Hyperoartia species we detected was the European river lamprey (*Lampetra fluviatilis*) with 0.1% of all reads. The mammal species with the highest read abundance was the European beaver (*Castor fiber*) with 4%, while the mallard (*Anas platyrhynchos*) with 1% and the graylag goose (*Anser anser*) with 0.5% were the birds with highest read abundances. A total of 17% of the reads were not assigned to species level. We found that the number of PCR replicates was positively correlated (p < 0.05) with both reads (rho = 0.843) and OTUs (rho = 0.924) (figure S1). Furthermore, PCR replicates showed high similarity values of shared OTUs across all samples, ranging from 85.53% to 97.1% (figure S2).

**Figure 2:**
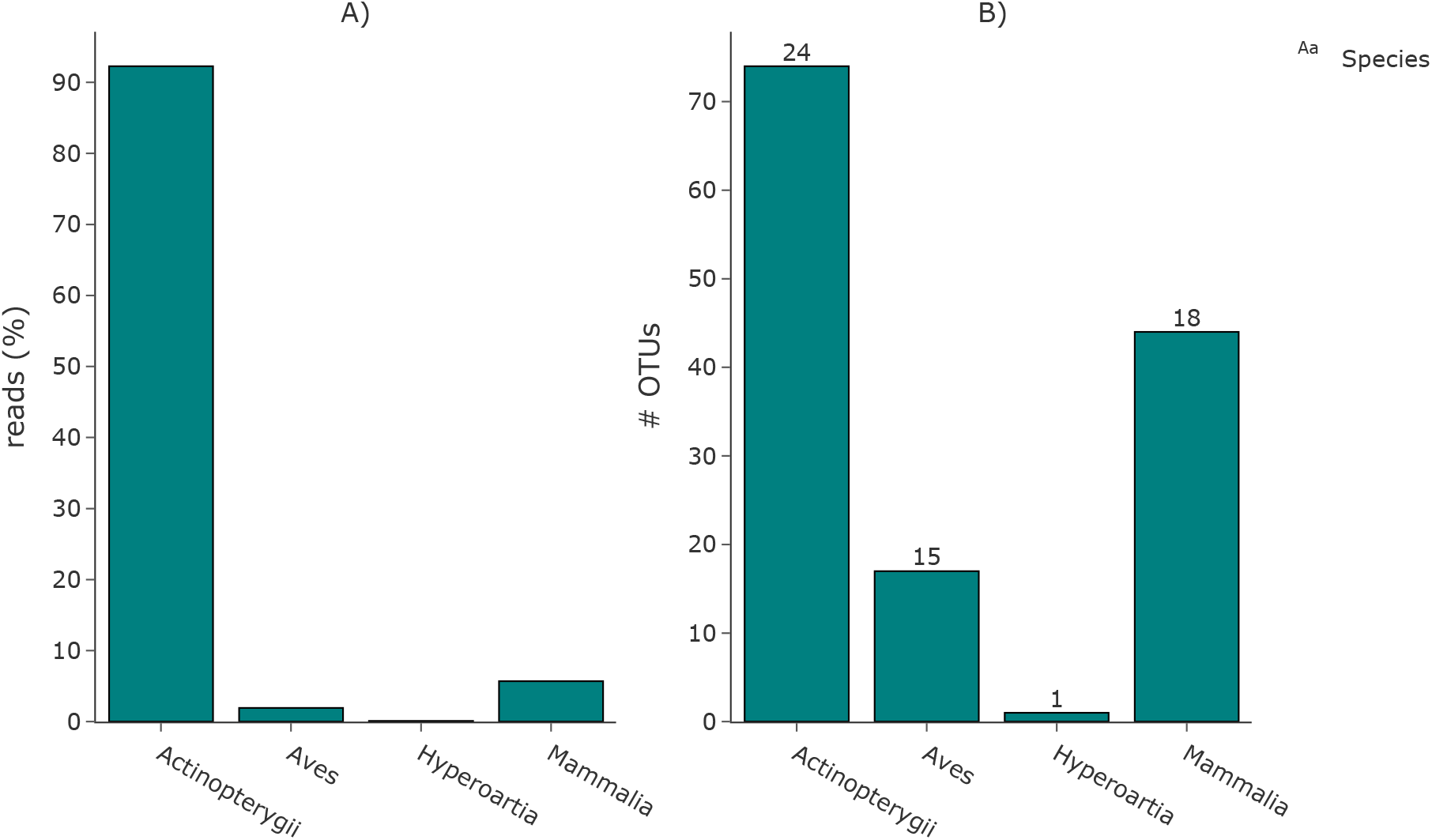
A) Percentage of reads assigned to the classes of Actinopterygii (ray-finned fish), Aves (birds), Hyperoartia (lampreys), and Mammalia (mammals). B) Number of OTUs assigned to the four classes. The number of assigned species is shown above the respective plot.

**Table 1:**
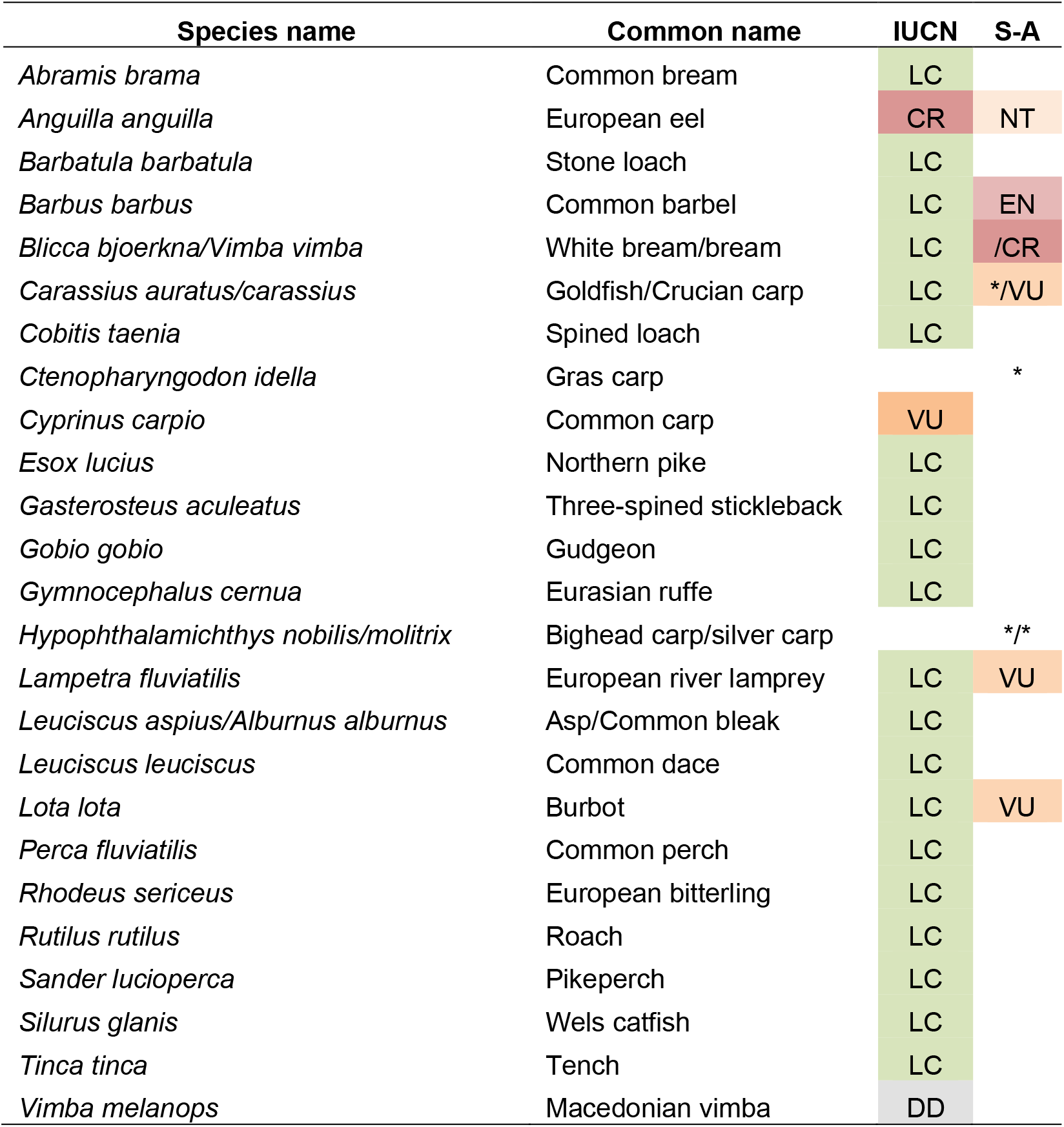
List of detected fish/lamprey species. The IUCN status and protection status of Saxony-Anhalt (S-A) are highlighted. Non-native species are marked with an asterisk.

| Species name | Common name | IUCN | S-A |
| --- | --- | --- | --- |
| *Abramis brama* | Common bream | LC |  |
| *Anguilla anguilla* | European eel | CR | NT |
| *Barbatula barbatula* | Stone loach | LC |  |
| *Barbus barbus* | Common barbel | LC | EN |
| *Blicca bjoerkna/Vimba vimba* | White bream/bream | LC | /CR |
| *Carassius auratus/carassius* | Goldfish/Crucian carp | LC | */VU |
| *Cobitis taenia* | Spined loach | LC |  |
| *Ctenopharyngodon idella* | Grass carp |  | * |
| *Cyprinus carpio* | Common carp | VU |  |
| *Esox lucius* | Northern pike | LC |  |
| *Gasterosteus aculeatus* | Three-spined stickleback | LC |  |
| *Gobio gobio* | Gudgeon | LC |  |
| *Gymnocephalus cernua* | Eurasian ruffe | LC |  |
| *Hypophthalmichthys nobilis/molitrix* | Bighead carp/silver carp |  | */* |
| *Lampetra fluviatilis* | European river lamprey | LC | VU |
| *Leuciscus aspius/Alburnus alburnus* | Asp/Common bleak | LC |  |
| *Leuciscus leuciscus* | Common dace | LC |  |
| *Lota lota* | Burbot | LC | VU |
| *Perca fluviatilis* | Common perch | LC |  |
| *Rhodeus sericeus* | European bitterling | LC |  |
| *Rutilus rutilus* | Roach | LC |  |
| *Sander lucioperca* | Pikeperch | LC |  |
| *Silurus glanis* | Wels catfish | LC |  |
| *Tinca tinca* | Tench | LC |  |
| *Vimba melanops* | Macedonian vimba | DD |  |

**Table 2:**
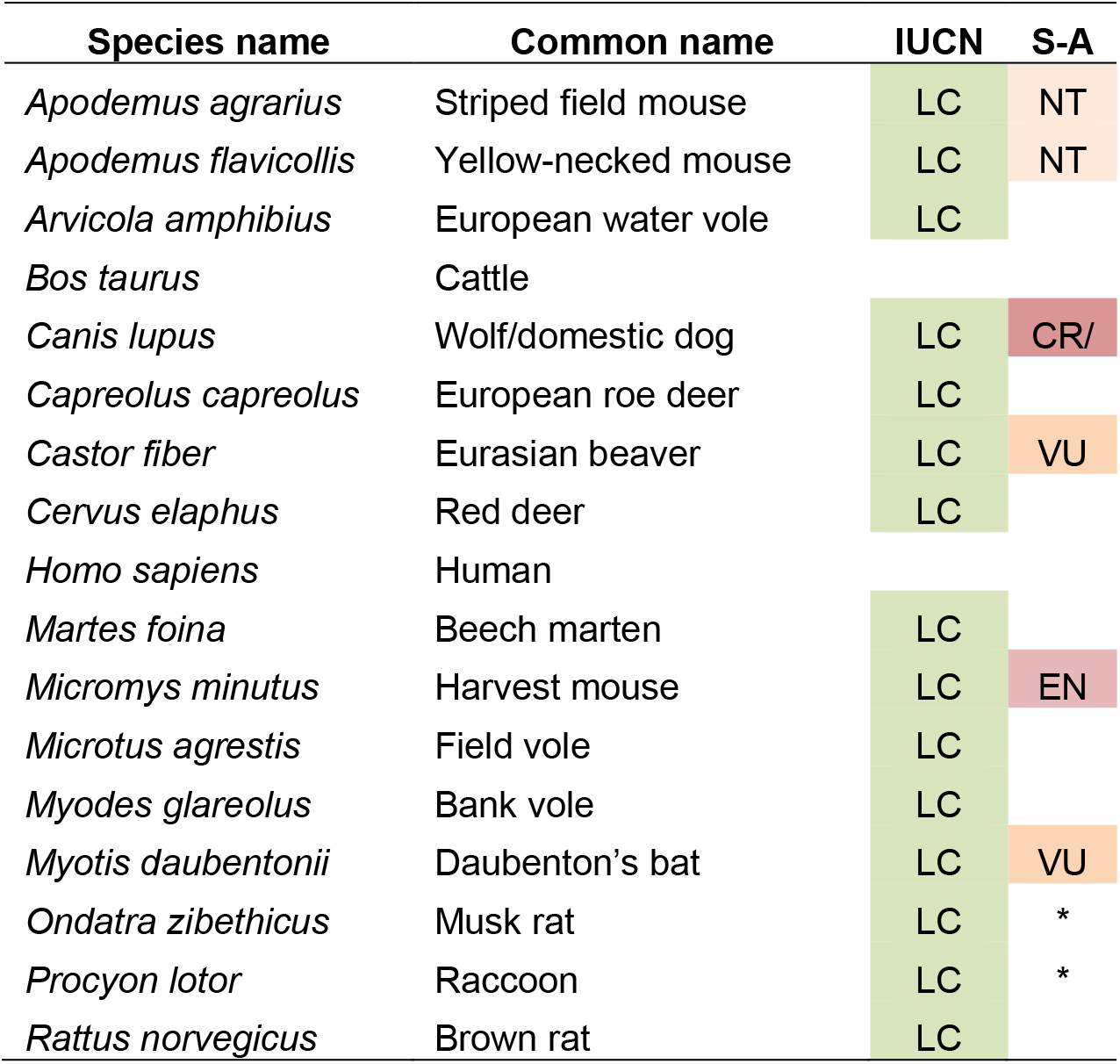
List of detected mammal species. The IUCN status and protection status of Saxony-Anhalt (S-A) are highlighted. Non-native species are marked with an asterisk.

| Species name | Common name | IUCN | S-A |
| --- | --- | --- | --- |
| *Apodemus agrarius* | Striped field mouse | LC | NT |
| *Apodemus flavicollis* | Yellow-necked mouse | LC | NT |
| *Arvicola amphibius* | European water vole | LC |  |
| *Bos taurus* | Cattle |  |  |
| *Canis lupus* | Wolf/domestic dog | LC | CR/ |
| *Capreolus capreolus* | European roe deer | LC |  |
| *Castor fiber* | Eurasian beaver | LC | VU |
| *Cervus elaphus* | Red deer | LC |  |
| *Homo sapiens* | Human |  |  |
| *Martes foina* | Beech marten | LC |  |
| *Micromys minutus* | Harvest mouse | LC | EN |
| *Microtus agrestis* | Field vole | LC |  |
| *Myodes glareolus* | Bank vole | LC |  |
| *Myotis daubentonii* | Daubenton's bat | LC | VU |
| *Ondatra zibethicus* | Musk rat | LC | * |
| *Procyon lotor* | Raccoon | LC | * |
| *Rattus norvegicus* | Brown rat | LC |  |

**Table 3:**
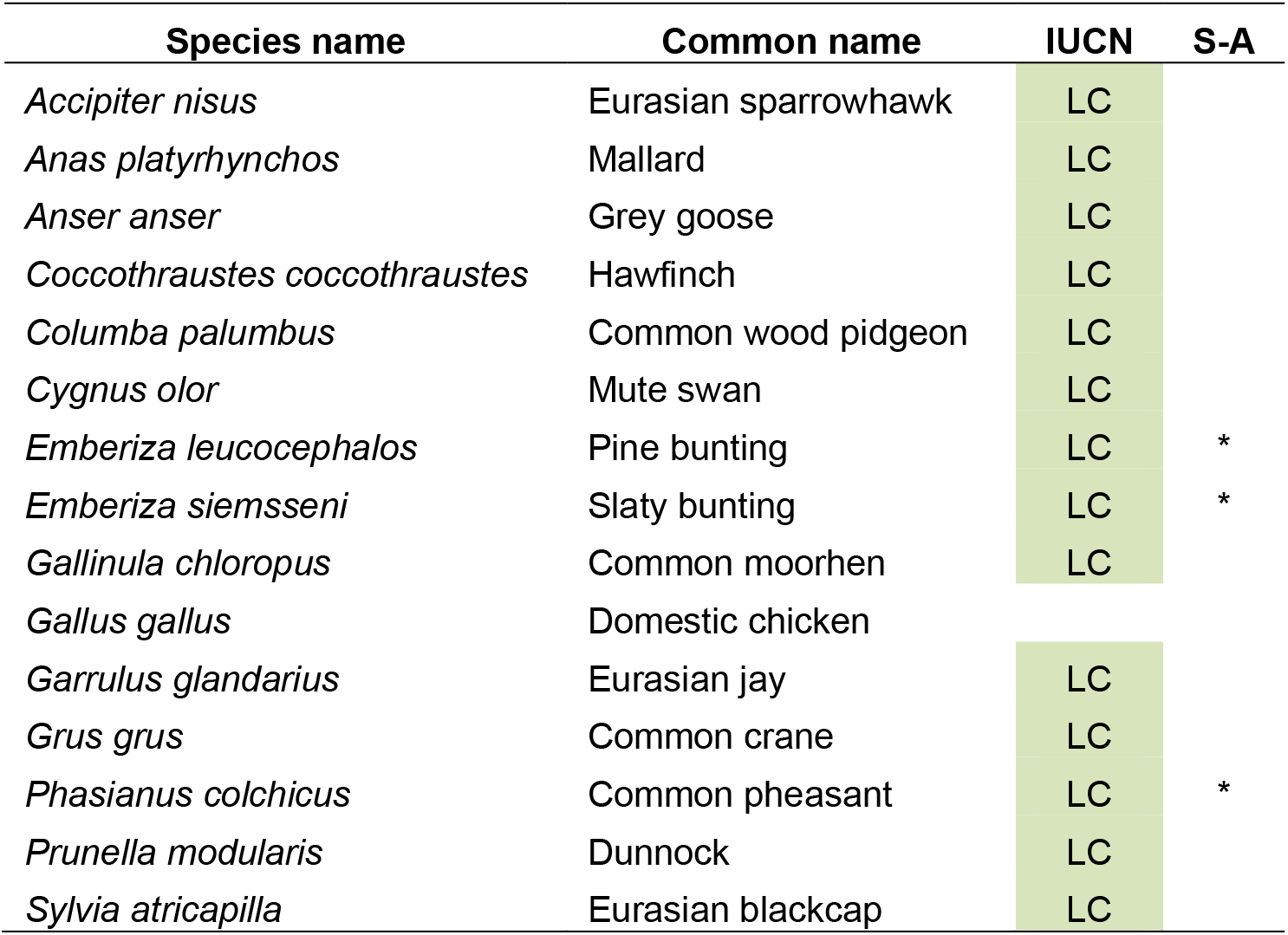
List of detected bird species. The IUCN status and protection status of Saxony-Anhalt (S-A) are highlighted. Non-native species are marked with an asterisk.

| Species name | Common name | IUCN | S-A |
| --- | --- | --- | --- |
| *Accipiter nisus* | Eurasian sparrowhawk | LC |  |
| *Anas platyrhynchos* | Mallard | LC |  |
| *Anser anser* | Grey goose | LC |  |
| *Coccothraustes coccothraustes* | Hawfinch | LC |  |
| *Columba palumbus* | Common wood pidgeon | LC |  |
| *Cygnus olor* | Mute swan | LC |  |
| *Emberiza leucocephalos* | Pine bunting | LC | * |
| *Emberiza siemsseni* | Slaty bunting | LC | * |
| *Gallinula chloropus* | Common moorhen | LC |  |
| *Gallus gallus* | Domestic chicken |  |  |
| *Garrulus glandarius* | Eurasian jay | LC |  |
| *Grus grus* | Common crane | LC |  |
| *Phasianus colchicus* | Common pheasant | LC | * |
| *Prunella modularis* | Dunnock | LC |  |
| *Sylvia atricapilla* | Eurasian blackcap | LC |  |

No consistent differences in the community composition between the field replicates along the 2 km stretch were found based on the NMDS results (dimensions=3; stress=0.75). Thus, we treated all samples as individual field replicates. To evaluate the effect of sampling effort on the detected species richness, we separately ran rarefaction analyses for fish/lamprey, mammals and birds (figure 3). Our results showed a substantial increase in detected species richness with increased sampling effort for all three groups. However, we observed a strong disproportionate increase between fish/lamprey species richness and mammal and bird species. Here, the fish/lamprey species showed the lowest increase from an average of 18.7 (±2.3 standard deviation) species in one sample to a maximum of 25 detected species in all 18 samples. The detected species richness of both mammals and birds increased substantially more. Here, we observed 5.7 (±1.7) mammal species and 3.4 (±1.5) bird species on average in one sample to a maximum of 18 and 15 species in all 18 samples. This accounts for an overall growth in detected species richness of 25.2% for fish/lamprey, 68.3% for mammals, and 77.3% for birds. The rarefaction curve for the fish/lamprey species showed its strongest increase in the inclusion of the first 8 replicates, accounting for 80% of the increase, and then nearing an asymptote towards the maximum number of 18 samples. The rarefaction curves of the mammal and bird species did not reach an asymptote but showed a consistent linear increase, indicating a further increase of species richness beyond the 18 samples. Overall, the majority of fish species (19 of 25) were detected in at least 50 % of the samples (figure 4). Only two fish species were solely detected in one sample (*Lota lota* and *Cobitis taenia*). As for the other vertebrates, the majority of species was detected in less than 50% of the samples, accounting for 13 of 15 bird species and 12 of 17 mammal species.

**Figure 3:**
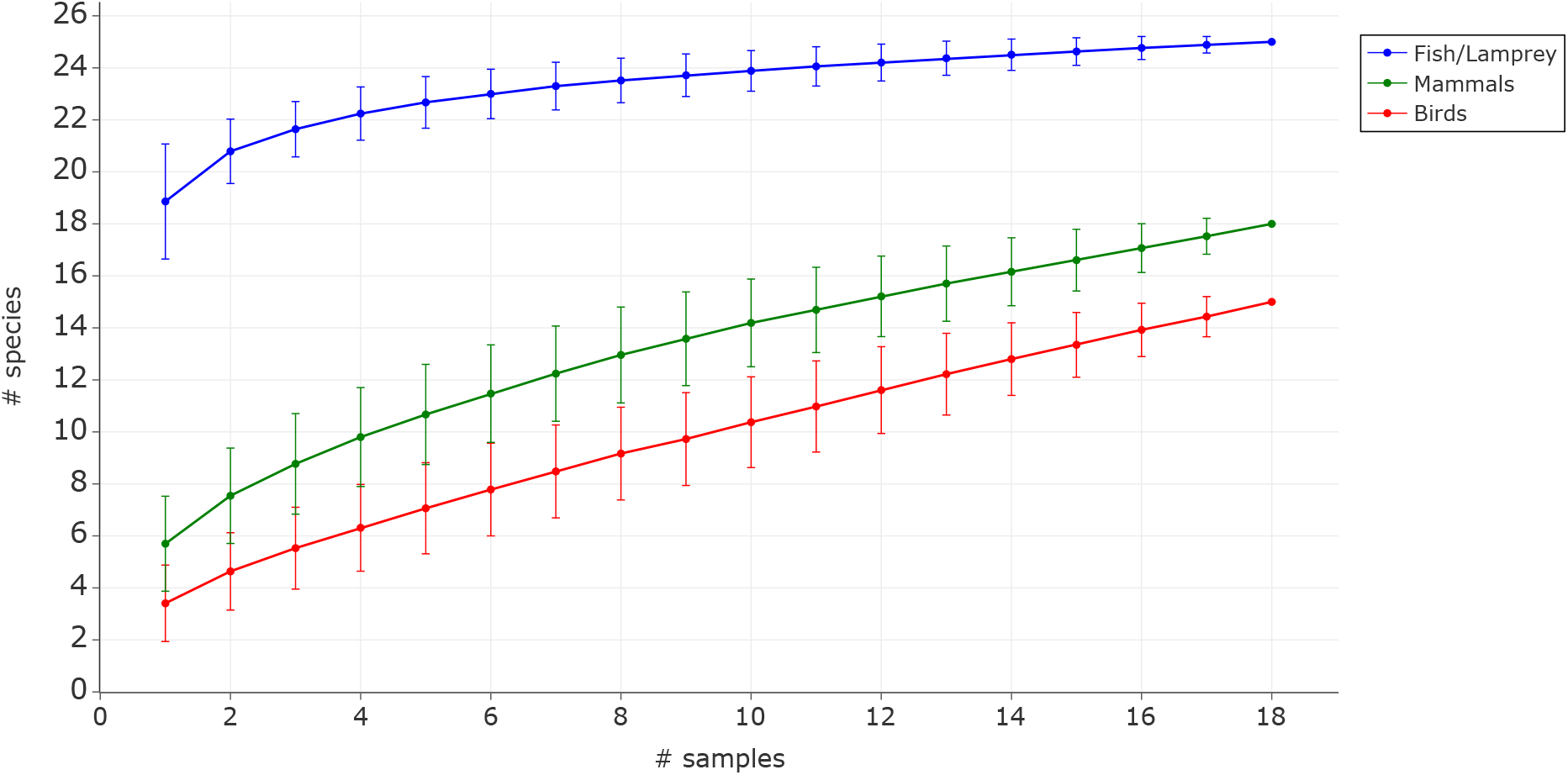
Rarefaction curves of the detected species richness of fish/lamprey (blue), mammals (green) and birds (red). Samples were randomly drawn 1000 times for each group to account for stochasticity.

**Figure 4:**
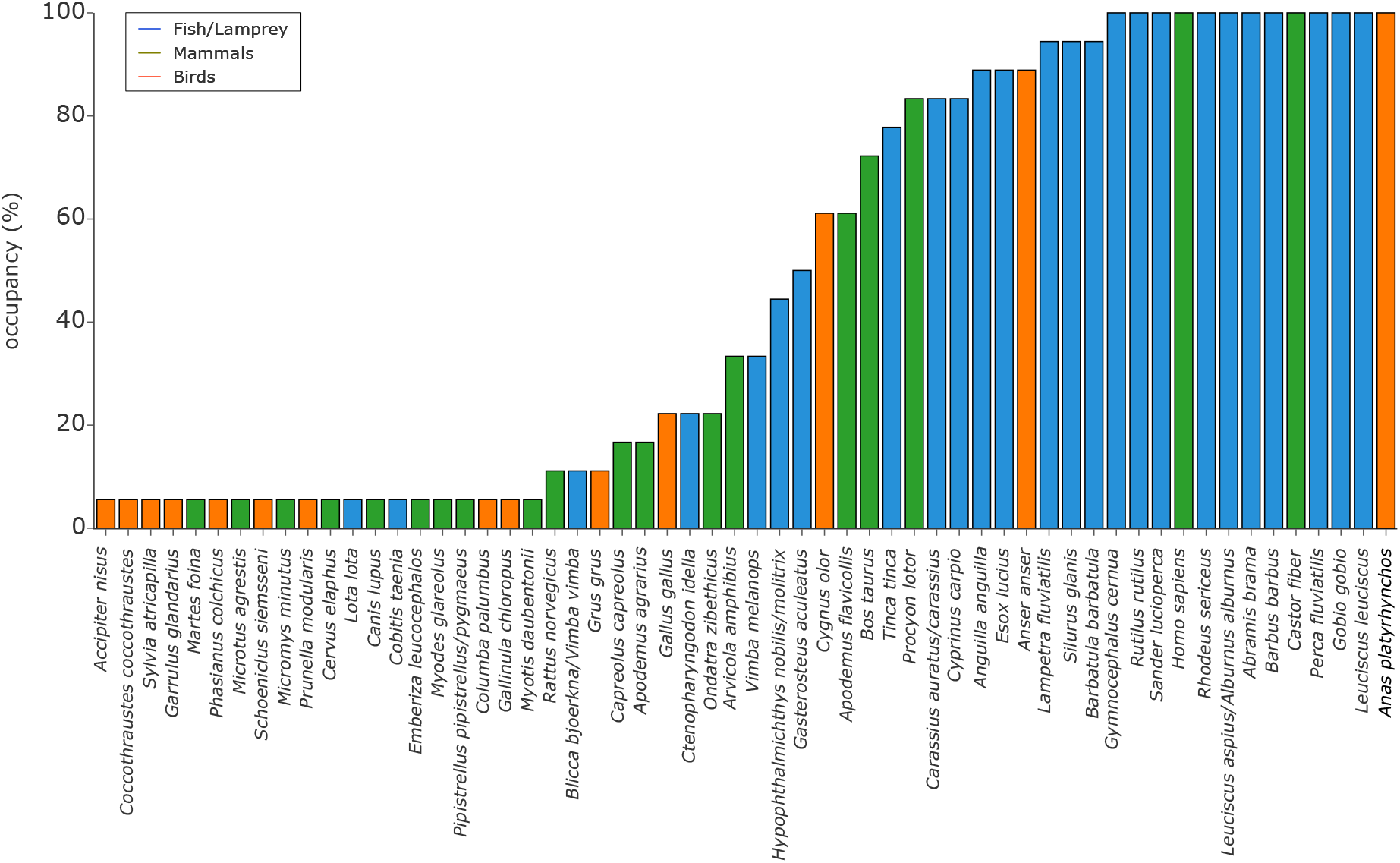
Occupancy of fish/lamprey, bird and mammal species across all samples.

## Discussion

### Detected fish biodiversity

Using eDNA metabarcoding, we successfully detected 25 fish species known to occur in the river Mulde and, further, even 50% of all fish species native to Saxony-Anhalt. Most fish species belonged to the order of Cypriniformes (66% of all species), which was expected since they are the dominant group in Central European rivers (Freyhof and Brooks 2011). The species that stood out in terms of read abundance (57.7% of all reads) was the common dace (*Leuciscus leuciscus*), followed by the Eurasian ruffe (*Gymnocephalus cernua*, 4%), and the common bream (*Abramis brama*, 4%). Quantitative interpretations of read counts and biomass or specimens abundance have been reported for fish (Bernd Hänfling et al. 2016; Ushio, Murakami, et al. 2018; Salter et al. 2019; Muri et al. 2020) but can be prone to several sources of bias. In our study, the sampling event took place during the spawning time of various fish species in spring. This can lead to a potential inflation of eDNA molecules of certain species that for example release their eggs and sperm into the open water, such as the common dace (Mills 1981). Furthermore, we cannot rule out primer specific bias that in- or deflates read counts of certain species. Thus, we here omitted correlations of read counts to specimen abundance or biomass and merely focused on species occurrence.

However, not all OTUs were successfully assigned to species level. We found multiple taxa where the 12S marker resolution was too low to distinguish between species and instead two species with identical similarity score were assigned. We manually checked these cases and found several OTUs for which both potential species were reported from the Mulde. For these ambiguous taxonomies we chose a strict approach and counted those cases as one entry. For example, we found the crucian carp and goldfish (*Carassius carassius* and *C*. *auratus*), where the crucian carp is the ancestry species of the domestic goldfish (Chen et al. 2020). Other closely related species we found are the white bream (*Blicca bjoerkna*) and vimba bream (*Vimba vimba*), the asp (*Leuciscus aspius*) and common bleak (*Alburnus alburnus*), and the invasive bighead and silver carp (*Hypophthalmichthys nobilis* and *H. molitrix*). Furthermore, the record of the Macedonian vimba (*Vimba melanops*) was puzzling, since it does not occur in Germany. We suggest that this hit resembles most likely a vimba bream (*Vimba vimba*), as we found this native species in our dataset and both are closely related (Hänfling et al. 2009), which may impact the taxonomic assignment. These findings confirmed that the tele02 marker is not suitable to distinguish all Central European fish at species level.

### Beyond fish eDNA metabarcoding: investigating bycatch detection

While most studies discard all non-target sequences (e.g., Evans et al. 2017; Li et al. 2018; Harper et al. 2019; Sales et al. 2020), we explicitly explored the legitimacy of the detected species and discuss whether they can inform other biomonitoring or species conservation activities. We here used the teleo2 primer pair that is known to amplify DNA of other vertebrate species than fish (Mariani et al. 2021). First, we could show that many vertebrate species besides fish were found as bycatch in our samples. We were able to detect a notable 22.2% and 7.4% of the whole native mammal and breeding bird fauna reported from Saxony-Anhalt, respectively. While only a minority of the detected species are water-bound or semi-aquatic, the majority inhabit agricultural, forest, and urban habitats, which accompany large parts of the upstream areas of the river Mulde. All organisms depend on water as a drinking source, which makes streams a sink for eDNA signals, transporting them downstream. The most represented group of the vertebrate species bycatch in terms of read proportions and species richness were mammals. The most represented order within the mammals was rodents (Rodentia). Here, high read proportions were assigned to the semi-aquatic Eurasian beaver (*Castor fiber*), which is reported to inhabit the river Mulde (German national FFH report, 2019). Furthermore, several terrestrial rodents were found, which often inhabit agricultural, and urban environments, such as the striped field mouse (*Apodermus agrarius*) or the Eurasian harvest mouse (*Micromys minutus*). Four species of even-toed ungulates (Artiodactyla) were detected: Cattles (*Bos taurus*) are livestock and graze on fields near the river. Roe deer (*Capreolus capreolus*) and red deer (*Cervus elaphus*) are known to be good swimmers and can easily cross rivers to reach new feeding grounds and thus release traces into the water. Three carnivora species were detected with eDNA. The putative detection of *Canis lupus* is most likely explained by the detection of domestic dogs (*Canis lupus familiaris*), which cannot be distinguished from one another based in the analyzed 12S region. However, since wolf populations have significantly increased over the last decades in central Europe (Chapron et al. 2014) and wolves have been reported from the area of the sampling site (LAU Saxony-Anhalt wolf observation report 2020; J. Arle pers. obs.), a detection of a wild wolf cannot be excluded. The two other detected carnivore species are the beech marten (*Martes foina*), a generalist and adaptable species inhabiting open areas and forests, and the invasive raccoon (*Procyon lotor*), which inhabits forests or urban areas and is a good swimmer that prefers freshwater associated habitats. Furthermore, three bat species were found, i.e., the Daubenton’s bat (*Myotis daubentonii*) and a pipistrelle species (either *Pipistrellus pipistrellus* or *P*. *pygmaeus*). The Daubenton’s bat relies on clean streams or lakes and hunts insects directly over the water surface (Vesterinen et al. 2013; 2016), which makes it very likely to introduce DNA traces into the water by dropping hair, skin, saliva, urine, and feces. The two detected pipistrelle species are closely related and were not distinguishable with the 12S marker. Birds, however, were generally less represented in both read proportions and species richness compared to mammals. An initial observation was that the detected birds are rather common species that occur in high numbers in the area, compared to the detected mammals. Several aquatic and marsh birds were detected, such as the mallard (*Anas platyrhynchos*), the graylag goose (*Anser anser*), the mute swan (*Cygnus olor*), and the common crane (*Grus grus*). While the first three species are common waterfowl in Germany all year round, the common crane is a migratory bird. Its detection falls directly in the spring migration, when large flocks of common cranes travel northwards, which makes a detection with eDNA very likely. Besides the waterfowl, most detected species belonged to passerine birds. Two puzzling species were detected that are not present in Germany: the pine bunting (*Emberiza leucocephalos*) and the slaty bunting (*Schoeniclus siemsseni*). Here, the most likely explanation is that the 12S marker is not suitable to identify them at species level and distinguish them from other bunting species that are inhabiting Germany and are abundant in the area, such as the common reed bunting (*Emberiza schoeniclus*) or the yellowhammer (*Emberiza citrinella*). Furthermore, no amphibians or reptiles were found in the dataset. Particularly the absence of amphibians was notable since at least frog species of the genus *Rana* and toad species of the genus *Bufo* are commonly occurring in streams and ponds in Central Europe. Since the detection of amphibians is possible with the here used tele02 primer (Mariani et al. 2021) the river Mulde is most likely not a suitable habitat for amphibians, especially during the reproductive season, which falls into the time of the sampling event.

### Streams as eDNA conveyor belts

We found no effect of sampling distance on fish species detection. Thus, although samples were collected at five distinct locations of the river Mulde, the 18 collected samples can be considered as individual field replicates rather than 2-4 specific replicates of 5 sites. The lack of a spatial signal is, on the one hand, not unexpected considering that sampling sites were max. 2 km apart, which is well in the range of reported transport distances of eDNA (Deiner and Altermatt 2014; Shogren et al. 2017; Nukazawa, Hamasuna, and Suzuki 2018). On the other hand source, state, transport, and fate of eDNA is anything but simple (Barnes and Turner 2016). While eDNA molecules are transported downstream in general, they are influenced by shedding, retention, and resuspension processes along the way (Shogren et al. 2017). Also, location and density of populations thus can greatly influence the detectability (Carraro, Stauffer, and Altermatt (2021). Community signal inferred via eDNA can thus be very site-specific (Cantera et al. 2021). Besides the spatial aspects, sampling time may be even more important in streams, as eDNA concentration can be increased for several taxa due to e.g., seasonal events such as spawning (Wacker et al. 2019) and migration (Thalinger et al. 2019). Also, water discharge drastically differs among seasons thus leading to different baseline concentrations, suggesting the use of hydrological models in eDNA assessments to increase reliability (Carraro et al. 2018).

### Disproportionate increase of fish and bycatch detection

Generally, the probability of detecting target DNA when present, i.e., the sensitivity of a method, depends on the concentration and dispersion of target DNA molecules at a site, the sampling design, and the laboratory workflow (Furlan et al. 2016). In previous studies both increased water volume filtered and implementation of field and PCR replicates were found to enhance the sensitivity of eDNA monitoring approaches (Civade et al. 2016; Bernd Hänfling et al. 2016; Evans et al. 2017; Beentjes et al. 2019). For example, a previous study on tropical stream fish species suggested that a saturation of tropical stream fish species detection can be reached when sampling 34 to 68 liters per site (Cantera et al. 2019).

Our results based on 18 1-L water samples showed that the detection probability of eDNA for non-fish vertebrate species differed substantially among samples. Comparing field samples, we found that fish species richness increased only by 25.2% when considering one versus all 18 samples. This was different for the detection probability of the non-fish bycatch vertebrate species. For mammals and birds, it increased by 69.3% and 77.3%, respectively, when including 18 field samples. While in aquatic organisms such as fish release all their DNA into the surrounding water, only traces of the predominantly terrestrial or aerial bycatch species enter the water leading to a much lower concentration. This expectation is also met when comparing eDNA detection of semi-aquatic bird and mammal species, which were detected in more than 50% of the samples (e.g., mute swan, graylag goose, and Eurasian beaver). Similarly high detection rates, however, were also found for domestic animals living in high abundances in the riparian area of the river (e.g., cattle). Harper et al. (2019) identified similar patterns that terrestrial mammal eDNA signals are weaker and can be detected less frequently than signals from semi-aquatic mammals, using a vertebrate specific primer (Riaz et al. 2011). We also found a high number of human reads in the samples, which are expected in an eDNA assessment in urban environments from various potential sources. Importantly, however, our negative controls did not show many human reads (in our case, a maximum of 4 reads in processed libraries) rendering lab contamination as unlikely. The human traces are likely derived from the original water sample. Nevertheless, low read counts are commonly observed in PCR negative controls and might also originate from laboratory contamination.

### eDNA bycatch: unexplored potential for conservation management

While often left aside in studies that focus only on fish biomonitoring, the relevance of detected non-fish bycatch species can be high. This holds true in particular for endangered or protected species that are often difficult to monitor and rely on sighting reports or intensive survey campaigns. Early reports of invasive species occurrence can also trigger timely management options. For the target taxa, i.e., fish, six of the 25 detected fish/lamprey species are listed as near threatened (European eel), vulnerable (crucian carp, European river lamprey, and burbot), endangered (common barbel), and critically endangered (bream) in the German federal state of Saxony-Anhalt. In the bycatch eDNA data, however, our results detected several mammal species that are classified as protected in Saxony-Anhalt, such as the striped field mouse and yellow-necked mouse (both near threatened), the European beaver and the Daubenton’s bat (both vulnerable), the Eurasian harvest mouse (endangered), and possibly the wolf (critically endangered). Although we were able to detect these endangered species, our findings only provide small insights into the whole vertebrate community, since this study was limited in terms of time coverage (one sampling event) and spatial coverage (2 km stretch of one river). The rarefaction analysis results predicted the detection of more mammal and bird species if more samples were collected. However, it is expected that advances in the standardization and operation of fish eDNA metabarcoding will lead to a higher rate of application in research and regulatory monitoring campaigns in the future. This goes in hand with an increasing amount of available bycatch data that can be analyzed and utilized. With hundreds or thousands of eDNA water samples that are potentially collected each year in countries that apply a nationwide routine monitoring, the coverage of water bodies and different habitats will automatically increase. This opens access to obtain highly resolved spatial and temporal data not only on fish distributions, but also detection patterns of bycatch species. The obtained data could be directly collected in online biodiversity databases and used for more comprehensive insights into vertebrate species occurrence and distribution. The additionally acquired data would then also be available for conservation planning and management and could help to increase the extent and accuracy of regional red lists and lead to a better intercalibration with the international red list. This accounts particularly for conservation monitoring under the EU birds directive (Directive 2009/147/EC, 2009), the “EU Regulation 1143/2014 on Invasive Alien Species” or the EU habitats directive (Council Directive 92/43/EEC, 1992), where data is generally hard to obtain and striking deficits in the monitoring coverage are known. For example, data on distribution and population sizes of the bird fauna is available in great detail, but observations are often conducted on a non-standardized, voluntary basis. For mammals, however, routine monitoring campaigns are even more scarce, since they are costly and time consuming. Here, the fish eDNA metabarcoding data could provide a notable increase of data points that can be sampled and analyzed under standardized conditions and can be evaluated by experts. The potential of obtaining new, additional information on terrestrial species, in particular elusive, rare or protected species without additional costs is immense and may also stimulate major international conservation initiatives currently developed in the context of the post-2020 CBD-framework.

Nevertheless, the reports of non-target species from fish eDNA metabarcoding have to be interpreted with particular caution. Environmental DNA metabarcoding comes with several challenges that can lead to both false negative and false positive identifications (Barnes and Turner 2016). This accounts particularly for terrestrial and aerial species, which are only temporarily interacting with the water and leave only marginal traces.

We also detected species that are generally unlikely to inhabit the catchment and thus likely represent a false-positive result. Here, potential sources are that the marker resolution is too low to distinguish species, the detected eDNA was already degraded or the reference sequences are incorrectly labeled. But also, introduction of eDNA via effluent from sewage plants or other influx can falsify the picture of the species distribution (Yamamoto et al. 2016). Particularly false positive signals must be flagged to avoid biased distribution patterns, when they can be identified as such. It also has to be considered that commonly used fish primers, such as the MiFish (Miya et al. 2015) and tele02 primers are optimized for fish and discriminate the amplification of other taxa, which will most-likely lead to a lower detection rate compared to fish species. To compensate for the primer bias, universal vertebrate primers (e.g., Riaz et al. 2011) could be used which would allow to monitor fish, mammal, bird, reptile, and amphibian species at once without additional sampling or laboratory efforts. However, if the main target of the routine biomonitoring remains to detect fish, specific primers might perform better. If the goal is to target groups other than fish, the additional usage of specific primer sets for mammals (MiMammal, Ushio et al. 2017) or birds (MiBird, Ushio et al. 2018) on the same DNA extract is possible, but would come with additional laboratory work and costs. These, however, are small and analysis can be automated (Buchner et al. in prep), thus the added value for other specific conservation and management programs can be immense.

## Outlook

Our results show that not only target fish but also bycatch species (i.e., birds, mammals) can be assessed reliably using eDNA metabarcoding. While the analysis of only few 1-L samples already delivered consistent estimates on fish species richness, the detected richness of non-target bycatch species steadily increased with the number of samples analyzed due to the lower concentration of eDNA molecules of these in the water. In total, we detected a notable 50% of fish species, 24% of mammal species and 7% of breeding bird species native to Saxony-Anhalt by sampling a single site at a single day only. In typical fish eDNA metabarcoding assessments, these bycatch data are typically left aside, yet, from a viewpoint of biodiversity monitoring they hold immense potential to inform about the presence of also (semi-)terrestrial species at the catchment site. Unlocking these data from the increasingly available fish eDNA metabarcoding information enables synergies among terrestrial and aquatic biomonitoring programs, adding further important information on species diversity in space and time. We thus encourage to exploit fish eDNA metabarcoding biomonitoring data to inform other conservation programs. For that purpose, however, it is essential that eDNA data is jointly stored and accessible for different biomonitoring campaigns, either at state, federal or international level.

## Supplement

**Figure S1:**
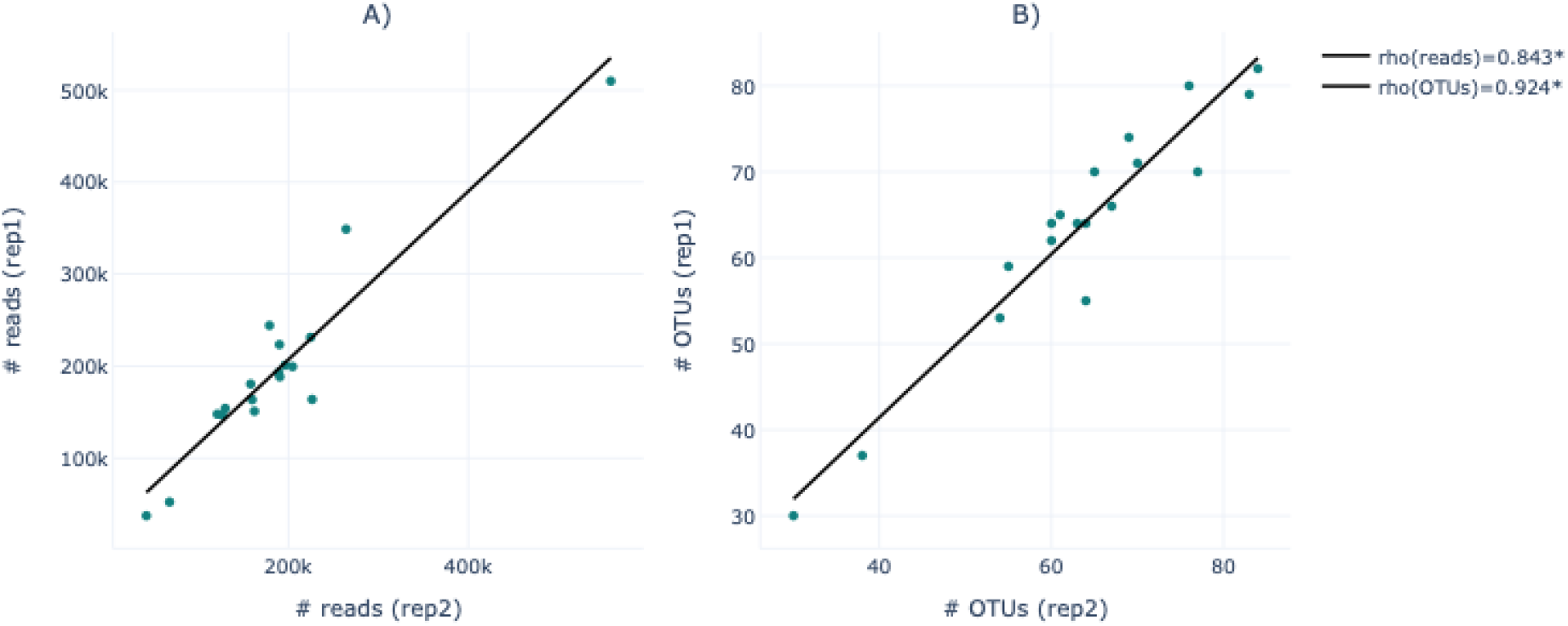
Spearman correlation analyses between 2nd-step PCR replicates for reads (A) and OTUs (B). Significant correlations (p ≤ 0.05) are marked with an asterisk.

**Figure S2:**
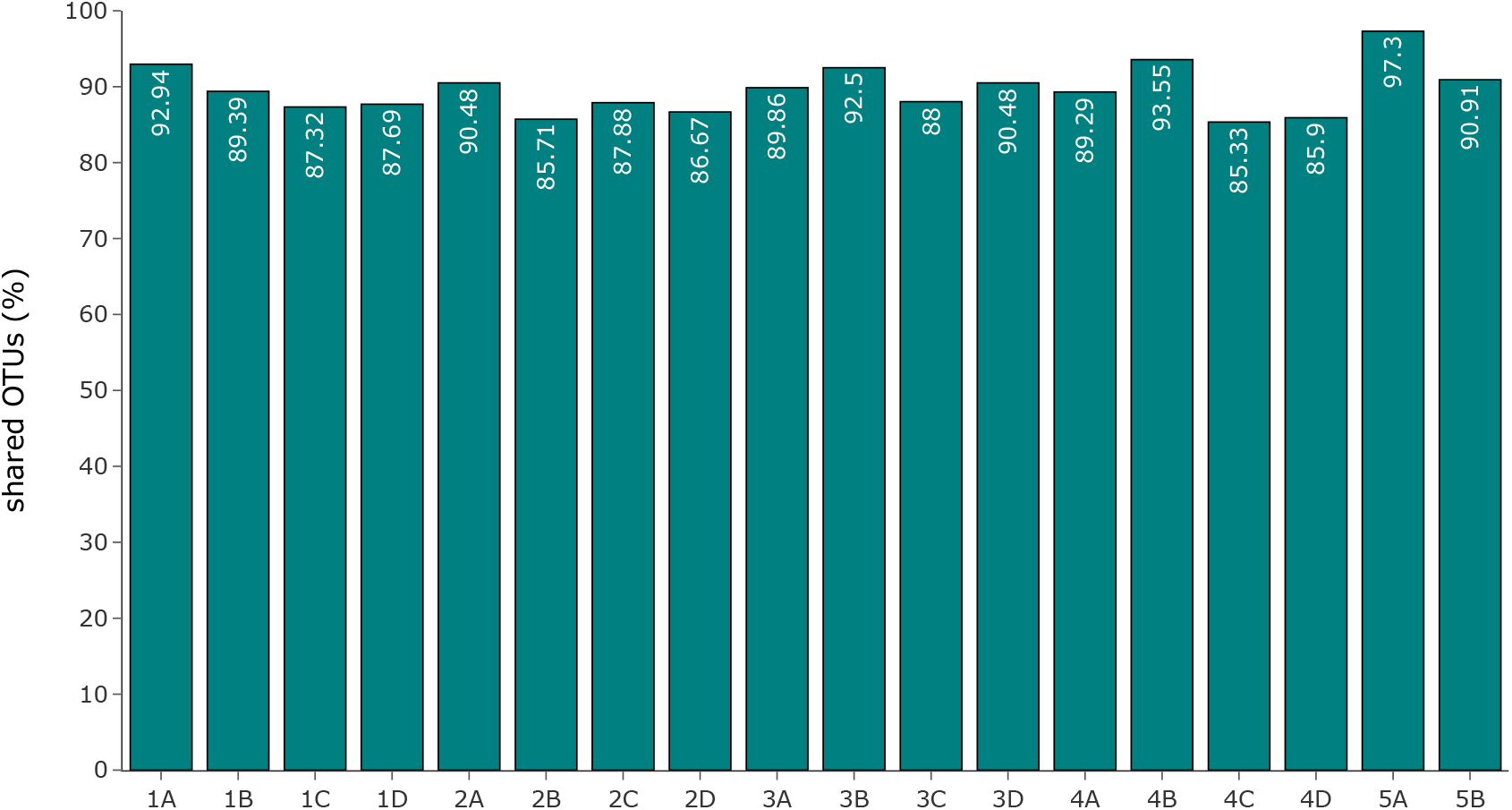
Proportion of shared OTUs between the two 2^nd^-step PCR replicates of each sample.

## Acknowledgements

We thank members of the Leese lab for comments and feedback on the study. We thank Falko Wagner (IGF Jena) for supporting the sample collection and discussions on the topic. This study was conducted as part of the GeDNA project, funded by the German Federal Environmental Agency (Umweltbundesamt, FKZ 3719 24 2040).

## Author Contributions

**Till-Hendrik Macher:** Conceptualization, Methodology, Formal analysis, Investigation, Visualization, Writing - original draft, Writing - review & editing. **Robin Schütz:** Formal analysis, Investigation, Writing - original draft, Writing - review & editing. **Jens Arle:** Resources, Validation, Writing - review & editing. **Arne J. Beermann:**Conceptualization, Formal analysis, Validation, Writing - review & editing **Jan Koschorreck:**Resources, Validation, Writing - review & editing **Florian Leese:** Conceptualization, Resources, Supervision, Project administration, Funding acquisition, Writing - review & editing.

